# DNA micro-disk for the management of DNA-based data storage with index and write-once-read-many (WORM) memory features

**DOI:** 10.1101/2020.04.22.054502

**Authors:** Yeongjae Choi, Hyung Jong Bae, Amos C. Lee, Hansol Choi, Daewon Lee, Taehoon Ryu, Jinwoo Hyun, Seojoo Kim, Hyeli Kim, Suk-Heung Song, Kibeom Kim, Wook Park, Sunghoon Kwon

## Abstract

DNA-based data storage has attracted attention because of its higher physical density of the data and longer retention time than those of conventional digital data storage^1–7^. However, previous DNA-based data storage lacked index features and the data quality of storage after a single access is not preserved, obstructing its industrial use. Here, we propose DNA micro-disks, quick response (QR)-coded micro-sized disks that harbour data-encoded DNA molecules for the efficient management of DNA-based data storage. We demonstrate the two major features that previous DNA-based data storage studies could not achieve. One feature is accessing data items efficiently by indexing the data-encoded DNA library. Another is achieving write-once-read-many (WORM) memory through the immobilization of DNA molecules on the disk and their enrichment through *in situ* DNA production. Through these features, the reliability of DNA-based data storage was increased by allowing multiple accession of data-encoded DNA without data loss.

---

DNA-based data storage, which encodes data to sequences of physical DNA molecules, was introduced as a solution to meet the exponentially increasing demand for cold data storage^1–8^. The two main advantages of DNA-based data storage are the high physical density of the data that allows for data storage of a building-scale backup centre in one laboratory tube^2,9^, and the centuries-long retention time of the data without electric power consumption^10,11^. Recently, the data-to-DNA encoding algorithms^1,2,4,6^ and DNA synthesis techniques^8^ were developed to establish error-less and cost-efficient DNA-based data storage. Based on these achievements, DNA-based data storage has been considered as an alternative to conventional data storage methods in the industry, especially for massive archival data storage.

To take the next step towards industrial application of DNA-based data storage, data management systems that are uniquely suited for access process of DNA-based data storage need to be developed. The conventional data storage media provides efficient search of the data from the data library and provides unlimited data access as write-once-read-many (WORM) or write-many-read-many (WMRM) memory. The current processes for DNA-based data access need to be improved in two features, multiple accession and indexing. First, multiple accesses of data in DNA-based data storage fundamentally disrupts the integrity of the original data because the process involves DNA enrichment techniques, such as PCR^4,5^ or hybridisation^12^. The DNA molecules that correspond to the original data are mixed with the reactant during the enrichment process; this results in an irregular abundance of distinct DNA molecules or bias. Multiple accession of data from the DNA library was previously attempted by amplifying the original DNA molecules, re-storing the data, and re-amplifying the amplified DNA molecules^4^. However, after multiple PCR reactions, PCR bias^13,14^ is randomly accumulated. Also, the bias is reflected in the next generation sequencing (NGS) result, in which particular population is missing and error rate is inevitably increased. To recover the data without error, the NGS coverage should be increased to obtain a population with low abundance. The current methods for multiple accession of data from the DNA library require adjustments in the NGS conditions. Consequently, this will increase the cost and duration of the data access. Moreover, because of the random and unpredictable nature of PCR bias^14^, PCR-based multiple accession will obstruct DNA-based data storage work flow, such as random access^5^ (see Supplementary Note 1 for details for discussions on the previous multiple reading process).

Second, proper indexing methods to support massive data management of DNA data storage are needed. To selectively retrieve a specific file from a massive data storage centre, metadata that compactly registers the names and sizes of each file should be used and accessed frequently on demand to search the data, instead of accessing the entire data storage. Recently, the random access^5,15^ scheme using DNA barcode indexing or, in other words, DNA barcode-based metadata that enables selective access of each file in the library has been developed to demonstrate the possibility of a DNA-based backup centre. In this system, the user should be provided with metadata that is stored in the DNA (e.g. the DNA barcode indices and names of files according to the barcode) to access a specific file in the entire DNA storage. Although the metadata is compact and very frequently accessed, the user needs to perform an additional DNA-based data access process including NGS on the metadata itself, in addition to the main NGS of the target file. While performing NGS on the massive target file is necessary and effective, NGS on the compact metadata is unnecessary, considering time and cost required for the NGS process. In addition to time and cost, the metadata is usually accessed much more frequently than the original data. Metadata based on DNA is vulnerable to disruption, since the data integrity deteriorate as the frequency of access increases. If a method with such an indexing system is to be developed, the metadata should be established in another data storage medium that allows frequent access.

To solve the two problems mentioned above, an efficient management method for DNA-based data storage, which enables multiple data accesses and indexing, needs to be developed. Here, we propose the DNA micro-disk, a QR-coded micro-sized disk that harbours data-encoded DNA molecules (Fig. 1). By introducing the simple process of immobilizing data-encoded DNA molecules on the optically indexed microparticle (Fig. 1a), we overcome the two intrinsic problems of DNA-based data storage. First, we established WORM memory in DNA-based data storage by immobilizing the data-encoded DNA molecules in a micro-disk. The immobilized DNA molecules on the disk are used as templates for in situ enzymatic reactions for DNA copy production (enrichment) (Fig. 1b). The DNA production cycle consists of chemical denaturation for the generation of single-stranded copies for data recovery, primer annealing, and elongation to make the double-stranded DNA linked to the polymer. Since the original DNA molecules with encoded data are immobilized on the micro-disk and copied on demand, the original DNA molecules are preserved rather than being corrupted. Second, we indexed and implemented optical metadata on the micro-disk by patterning the QR coded-substrate. The QR index, which describes the metadata, contains the primer sequences and names of the files that are essential for the enrichment of data-encoded molecules from the immobilized DNA molecules on the disks. Instead of going through an additional NGS process for metadata, the metadata encoded in the QR code can be accessed by simple imaging within a few seconds. We established a metadata searching system that not only allows frequent access, but is also orthogonal to the DNA-based data storage. By overcoming the two limitations with the DNA micro-disk, we further demonstrate that a single micro-disk can store multiple files, and each file is recovered selectively by producing multiple DNA copies from the original immobilized DNA molecules on the micro-disk. Even though the copies of DNA are consumed during data access, we observed that the original DNAs on the micro-disks are preserved with minor discrepancy. We believe that this simple-yet-powerful system will accelerate the practical use of DNA-based data storage.

**Figure 1.**
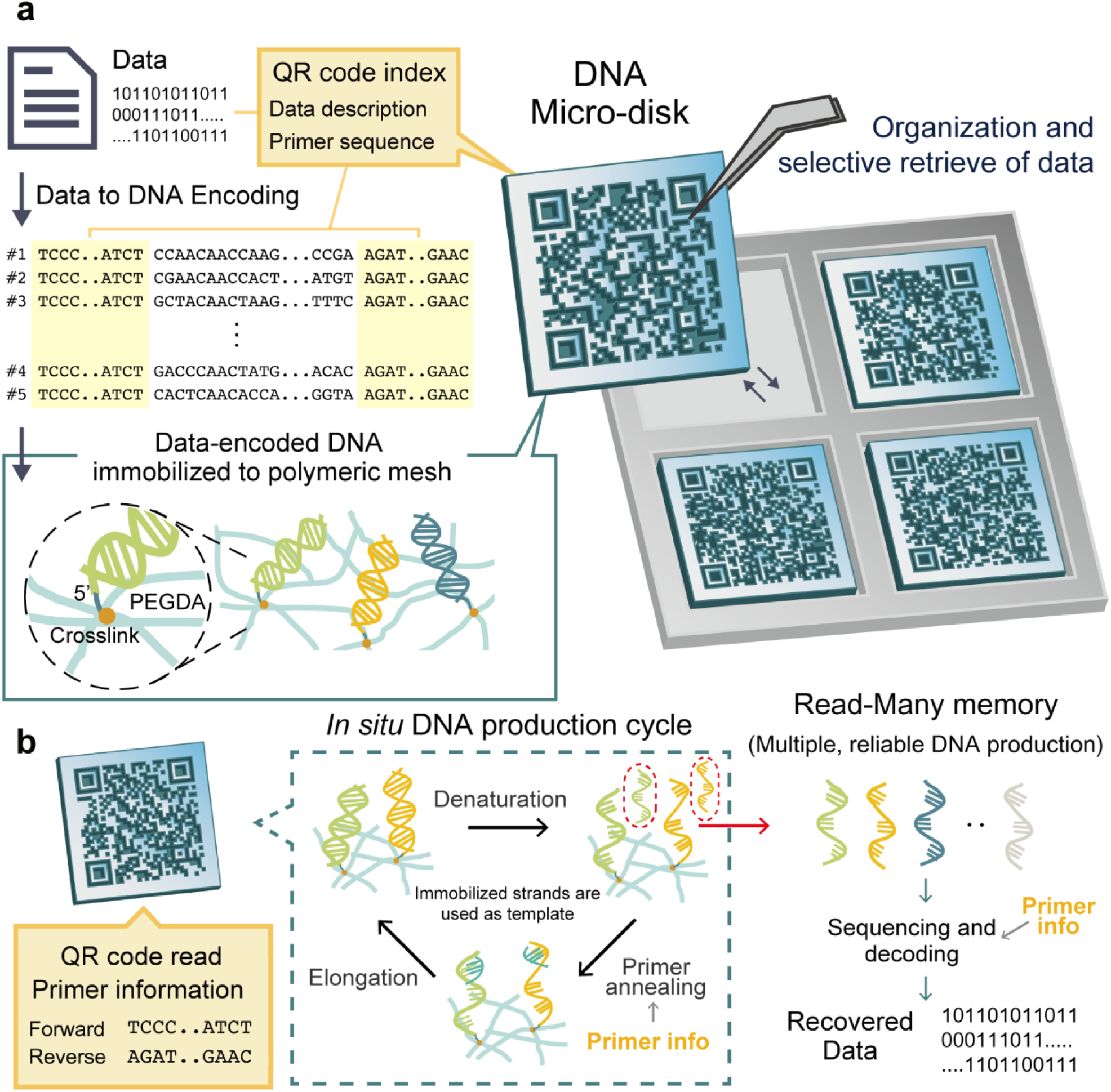
DNA micro-disk manages DNA-based data storage with index and write-once-read-many (WORM) memory features. **a**, The data-encoded DNA templates are immobilized on the polymeric mesh of the DNA micro-disk. Information of data-encoded DNA, such as primer information for amplification of the templates and data descriptions, is indexed in the QR code. DNA micro-disks can be organized on a microwell array and selected after QR code index scanning using micro-scale manipulation tools such as tweezers or pipettes. **b**, Data-encoded DNA copies from the DNA micro-disk are produced *in situ* through chemical denaturation of the DNA. Afterward, double-stranded DNA is restored by primer annealing and elongation. Through this process, immobilized DNA is preserved, and multiple accession of the data are enabled. DNA copies are used for sequencing and decoded for data recovery. The primer information described in the QR code is used for DNA production and sequencing.

DNA micro-disks of various sizes and data volumes were fabricated through optofluidic lithography^16–18^ and organized for increased accessibility of DNA-based data storage (Fig. 2, Supplementary Fig. 1). The data-encoding DNA library was amplified with a primer set, one of which was modified with acrydite, and mixed with photocurable polymer solution to make a pre-fabrication solution. A patterned ultraviolet (UV) light with a wavelength higher than 315 nm to minimize DNA damage was irradiated on the solution via a digital micromirror device to generate the micro-disk by photopolymerization. According to the UV pattern with intensity difference, the geometrically different (i.e. physical code) particles were fabricated (Figs. 2a, b). The physically engraved QR code on disk has the advantage of recovering the index even when the disk is physically damaged, due to the error-correction ability of the code^19^. The resolution of the system is 1 µm by the lithographical technique, and a micro-disk with a 100 × 100 µm cross-section could contain a version 20 QR code that can record 1249 alphanumeric characters. This not only is enough to contain the index information but also allows the recovery of the index information in case of micro-disk damage. We decided the size of the disk to be on the micro-scale. The micro-sized disk is simply manipulated and accessed with a mouse pipette and tweezers, and it can store a few gigabytes as a minimum storage capacity. Disk manipulation is important because while DNA is acquired through enzyme-mediated reactions in a solution, the disks must be handled individually. The mobile disks enable various forms of storage by following the demand. For example, even if disks containing different data are stored together without organization to minimize the volume of storage, the indexing is preserved, and the data do not interfere with each other. Additionally, the micro-disks can be assembled onto a microwell^20,21^ for index organization and imaging, increasing the accessibility (Fig. 2c).

**Figure 2.**
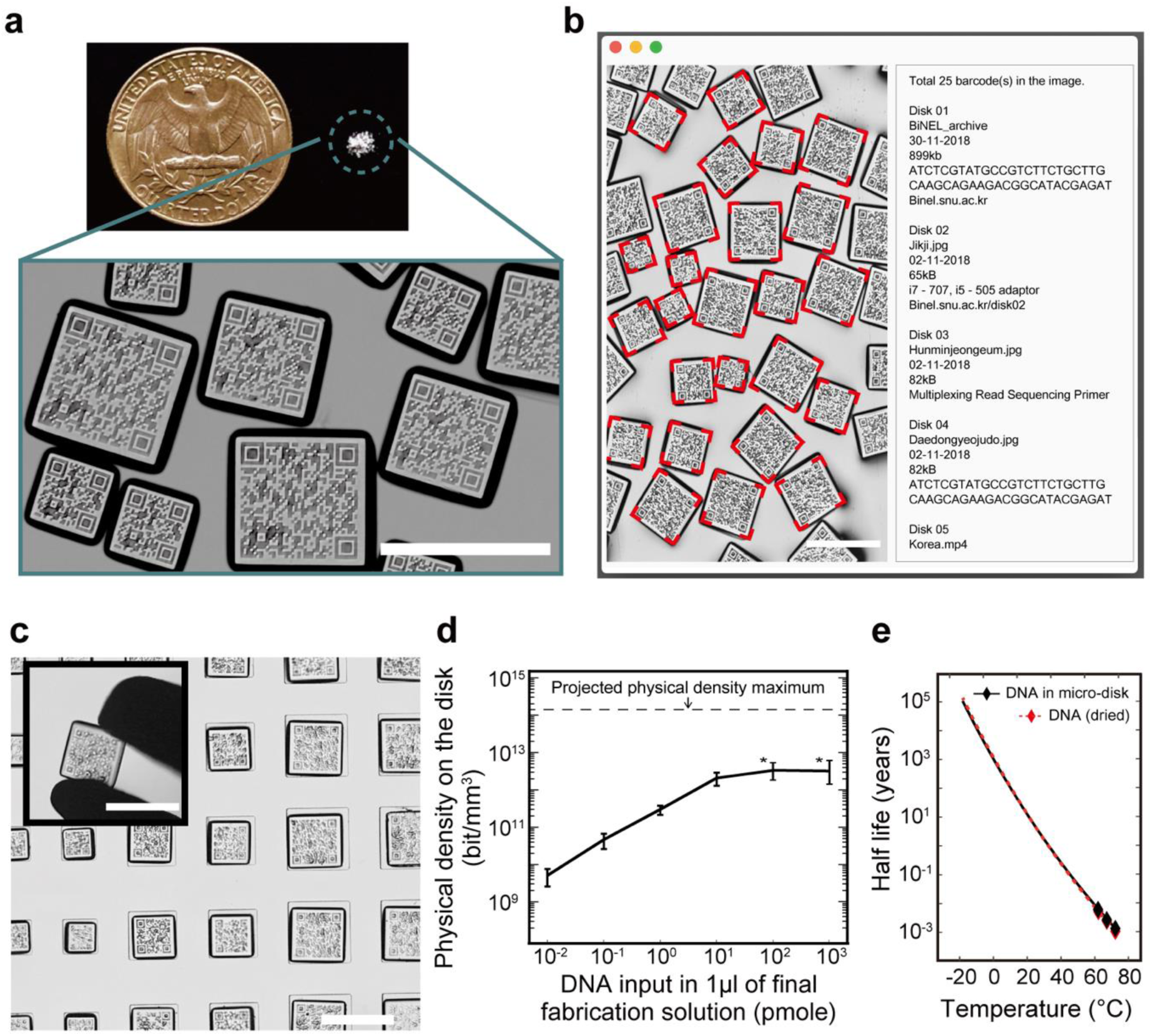
DNA micro-disk enables indexing and efficient access of data-encoded DNA libraries. **a**, DNA micro-disks of various sizes. Scale bar: 500 µm. **b**, QR codes of DNA micro-disks of various sizes are read for data access. Scale bar: 500 µm. **c**, DNA micro-disks are handled by tweezers or pipetting. DNA micro-disks of various size are assembled on the microwells. Scale bar: 500 µm. **d**, Depending on the DNA concentration in the pre-polymer solution during fabrication, micro-disks can be fabricated with various data capacities (n = 10). The error bars represent standard deviations. When the amount of input DNA exceeds 100 pmoles in 1 µl of solution, DNA cannot be completely dissociated in the fabrication solution and eventually precipitates (indicated by asterisks, Supplementary Fig. 2a). The projected physical density maximum describes the possible increment of physical density upon the application of the current state-of-the-art protocol on DNA-based data storage. The details regarding the calculation method are described in Supplementary Note 2. **e**, Extrapolated half-life of 160-bp DNA stored in dried condition or in a DNA micro-disk at 50% relative humidity. Dots indicate the value obtained from the experiment of DNA in a micro-disk (see details in Supplementary Note 3).

To quantify the number of molecules and data that can be incorporated into the micro-disks, we fabricated 250 x 250 x 50 µm micro-disks with different concentrations of 160 nucleotide-long DNA in the pre-fabrication solution (Fig. 2d, Supplementary Fig. 2a). As the concentration of DNA increased, the DNA molecules on the micro-disk also increased when quantified by qPCR (see Materials and Methods for details), and saturated at approximately 10^1^ pmol/mm^3^ (973 ng/mm^3^), as it reached the solubility limit of DNA in the pre-fabrication solution (Supplementary Fig. 2b). We achieved a physical density of approximately 10^12^ bit/ mm^3^ for single micro-disk. This density can be potentially increased to approximately 10^14^ bit/mm^3^ using the state-of-the-art technique of DNA-based data storage^22^ that can store 17 exabytes in single gram of DNA (see Supplementary Note 2 for the method used for calculation). Although the potential value of physical density of our proposed technique is three orders lower than the current state-of-the-art method^22^ and one order lower than the DNA fountain method^4^, we achieved a higher physical density (more than ten thousands) than those of conventional data storage methods, such as hard magnetic tape or flash memory^23^. Also, because micro-disks are mobile, physical density of multiple micro-disks is similar to that of single micro-disk, when the multiple micro-disks are stored in one vial, without organization.

Additionally, we extrapolated and compared the half-life of DNA micro-disk storage and dried DNA (or ambient DNA-based data storage) by performing an accelerated aging experiment at 50% relative humidity as a function of temperature (Fig. 2e, see Supplementary Note 3 for details). The durability of dried DNA and DNA stored in the micro-disk was comparable, higher than 100 years at temperatures lower than 10°C.

The DNA micro-disk allows for DNA-based data storage with WORM memory (Fig. 3). The micro-disk DNA can be accessed many times through the polymerase chain reaction (PCR), which enriches the data-encoded DNA templates linked to the polymer mesh. During the enrichment, there may be damage to the original template due to nuclease activity or heat during thermocycling. To minimize this damage, *in situ* DNA production was performed. Double-stranded DNA on the micro-disk was denatured at basic pH, and single-stranded DNA that detached from the disk was used for sequencing. After denaturation, primer annealing and elongation were conducted to restore the double-stranded DNA. We used this strategy to verify that the DNA copies were produced evenly and to confirm that the produced DNA retained the original characteristics in repeated copies. We fabricated a 250 × 250 × 50 µm micro-disk with 10^7^ DNA molecules that stored an image file of the Jikji (Supplementary Fig. 5), a Korean heritage item (Fig. 3a). During the 20 cycles of multiple reading, micro-disks produced similar amounts of DNA molecules. We were able to acquire the decay rate of the DNA molecules while multiple reading, by linearly fitting *log10*(molecule number produced) vs. cycle. The slope was −0.001 (R^2^ = 0.60) or, in other words, 0.23% of the DNA molecules was lost for each DNA production cycle. As a result, half-life of multiple reading is around 300 cycles. In addition to the molecule number, by sequencing the produced DNA copies and comparing them to the designed DNA fragment through NGS, we checked how each fragment was aligned. We confirmed that the calls per million reads in the 1^st^ cycle were similar to those in the 20^th^ cycle, and no designed fragment was lost in a million reads (Fig. 3b). In comparison, we experimentally analyse the conventional multiple readings of DNA-based data storage^4^. With fixed NGS coverage (200x) and physical density, we amplified the DNA molecules using PCR, read the data, stored 10^7^ DNA molecules, and re-amplified the stored DNA molecules (this is one reaction, Supplementary Fig. 6). After 20 reactions, approximately 20% of the fragments were not found, and therefore, the data could not be recovered in a million reads, representing a ~200× sequencing coverage. Therefore, WORM data storage was enabled, increasing the reliability of DNA-based data storage.

**Figure 3.**
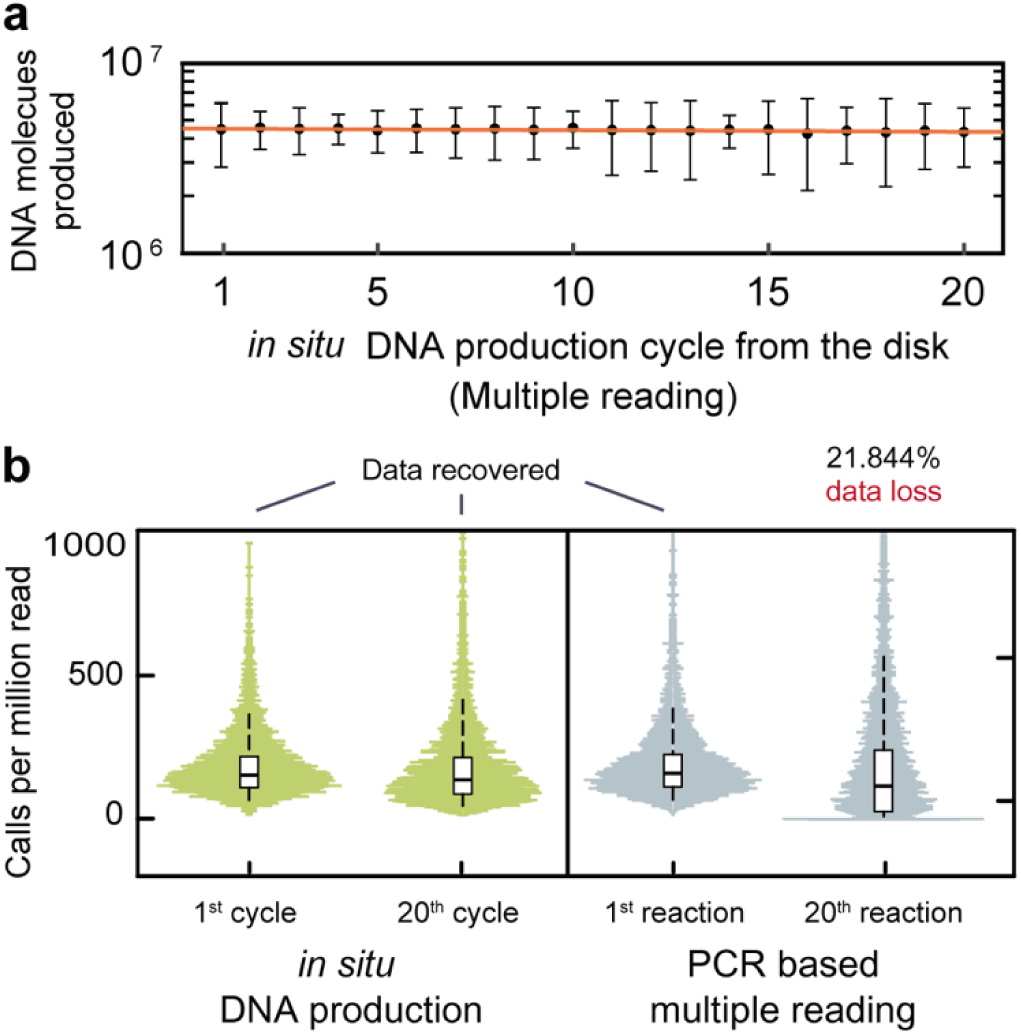
WORM memory DNA-based data storage is enabled through *in situ* DNA production from DNA micro-disks. **a**, Data-encoded DNA copies from the DNA micro-disks are produced by single elongation and chemical denaturation for 20 cycles and quantified (black, n = 20). The error bars represent standard deviations. A linear fit to the data (*log*_10_(molecules produced) vs. cycles) is shown in orange, R^2^ = 0.60. **b**, After 20 cycles of DNA production, the original characteristics such as the bias profile (produced by analysing the probability function of calls per million reads and the error rate) of the DNA copies from the micro-disk were preserved. In conventional DNA-based data storage, data are accessed multiple times by amplifying the original DNA and re-amplifying it (Supplementary Fig. 6). After 20 reactions, however, the bias profile and the error rate of the DNA template were not preserved, and data access was disabled, since the uncalled design was more than 20%. For each amplification cycle, 10^7^ molecules from the last cycle were amplified.

By applying the read-many feature to the DNA micro-disks, various files are stored on a single disk, and a specific file is selectively accessed multiple times (Fig. 4). In previous DNA-based data storage methods, if a user wanted to select and recover a specific file from an encoded DNA library of multiple files that were not physically separated, the targeted enrichment of DNA through designed primers or the random access^5^ was used. However, during the selection process, molecules that encode unselected files are corrupted without maintaining the original molecular population distribution (Supplementary Note 1). As a result, redundant copies of the DNA library are necessary as many as the number of the accesses desired. In contrast, when copies of DNA micro-disks are generated and used repeatedly for targeted amplification from the original template, the disadvantages of the random-access methods are eliminated. This is because there is no risk of losing the original template. We stored three different image files related to the Korean heritage, the Daedongyeojido (map), Jikji (document), and Hunminjeongum-Hyerye (document) (Supplementary Figs. 5,7,8), with different reverse primers on a single DNA micro-disk. We selectively recovered each file by multiple cycles of DNA production and the random access using the primer set information from the QR code.

**Figure 4.**
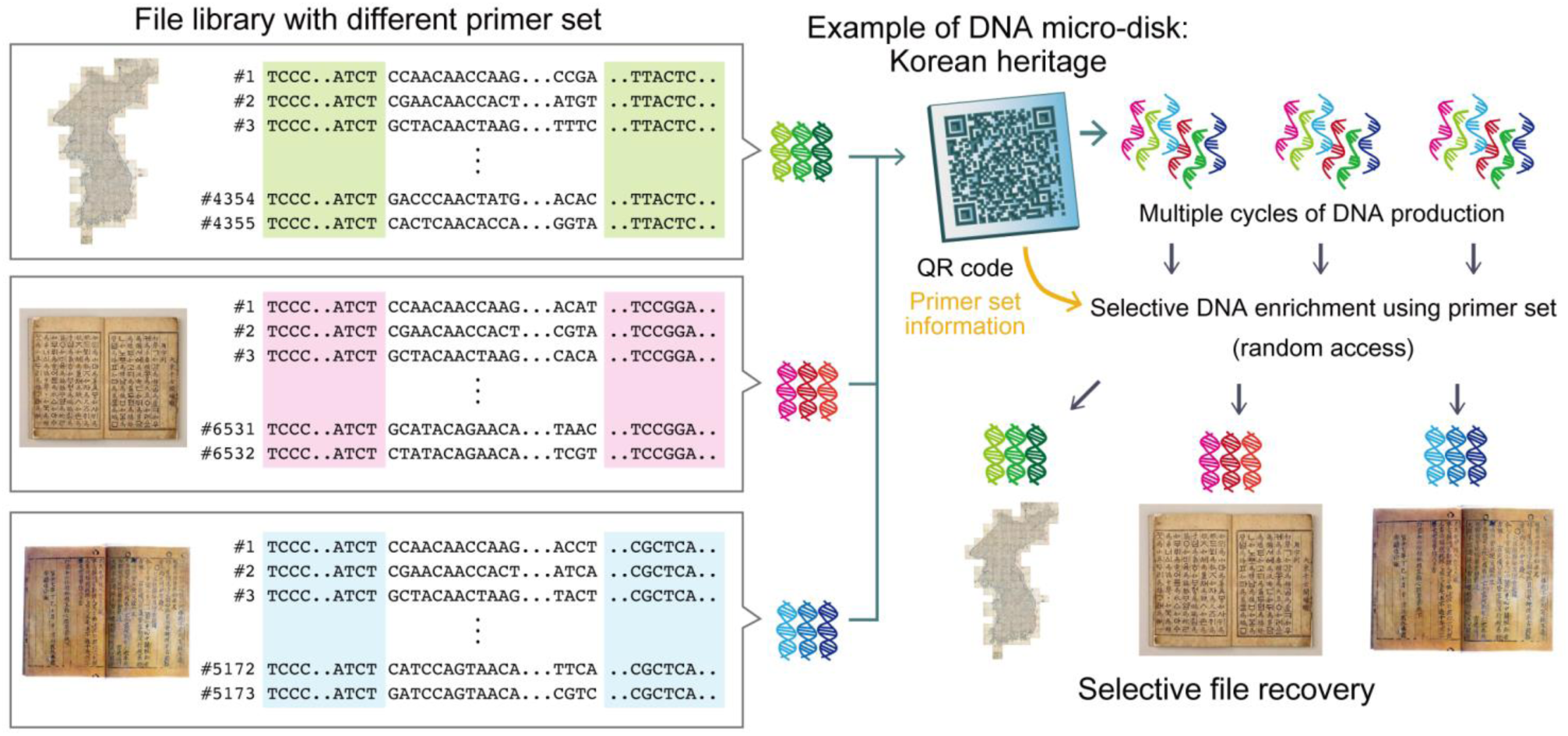
Single DNA micro-disk stores multiple files, and each file is recovered selectively through multiple DNA production and the random access^5^. Three different files from Korean heritage were encoded, synthesized as DNA with different primer sets, and stored in a DNA micro-disk. Whole DNA molecules were produced multiple times by *in situ* DNA production cycles. By utilizing the primer set information for each file, we selectively enriched the DNA and recovered the corresponding file. The image of the Hunminjeongum-Haerye was originally posted by the Cultural Heritage Administration of the Republic of Korea under the Korea Open Government License type 1.

In this letter, we propose the DNA micro-disk, which allows the management of DNA-based data storage in terms of accessibility and reliability (i.e. indexing and WORM). To our knowledge, the platform first introduces the metadata on data-encoded DNA molecules. Metadata is critical for overall system performance, especially in massive scale data storage systems^24,25^. The proposed platform has an additional advantage over the existing DNA management system based on DNA pellet microarray^15^ in preventing contamination between distinct DNA libraries by allowing immobilization of DNA on the micro-disk. In future, by optimizing the porosity of the polymer and the DNA crosslinking ratio, the physical density of DNA-based data storage can be further increased to the that of powder-based DNA based data storage. Finally, in the near future, with the development of an automated system that enables disk imaging and assembly, the DNA micro-disk system could be practically implemented as a data storage method.

## Acknowledgements

This work was partially supported by the Samsung Research Funding Center of Samsung Electronics under Project Number SRFC-IT1601-08, Global Research Development Center Program through the National Research Foundation of Korea (NRF) funded by the Ministry of Science and ICT (MSIT) (2015K1A4A3047345), Basic Science Research Program through the NRF funded by the Ministry of Education(NR F-2019R1I1A1A01059874), and grant from Kyung Hee University in 2015 (KHU-20150801). The Institute of Engineering Research at Seoul National University provided research facilities for this work.

## Author contributions

Yeongjae Choi, Hyung Jong Bae, Taehoon Ryu, Wook Park and Sunghoon Kwon initiated and designed the experiments. Yeongjae Choi, Hyung Jong Bae, Amos C. Lee, Wook Park and Sunghoon Kwon wrote the manuscript. Yeongjae Choi, Taehoon Ryu, Amos C. Lee, Hansol Choi, Jinwoo Hyun, Suk-Heung Song, Seoju Kim, and Hyeli Kim, Kibeom Kim conducted the research including DNA synthesis and analysis.

## Data availability

All sequencing data are available in Sequence Read Archive (SRA) under accession number PRJNA555140. All other data are available from the authors upon reasonable request.

