## Supplementary Information for "DNA micro-disk for the management of DNA-based data storage with index and write-once-read-many (WORM) memory features"

**This file includes:**

Supplementary Notes 1 to 3
Supplementary Figures 1 to 8

### Supplementary Note 1: Data quality change during the data access

The process of data access in DNA-based data storage requires dissociation of the DNA pellet as a whole and selective DNA enrichment. During DNA enrichment process, such as PCR[4,5] or hybridisation[12], The DNA molecules that correspond to the original data are mixed with the reactant during the enrichment process; this can result in an irregular abundance of distinct DNA molecules (PCR bias) or physical density lost.

i) PCR-based enrichment method

For PCR-based enrichment method, irregular abundance of distinct DNA molecules in the amplified population (or PCR bias[13,14]) is occurred after the reaction. Previously, Erlich and Zielinski[4] introduced the PCR-based multiple accesses of DNA-based data storage[4]. However, after the multiple PCR reaction, PCR bias is accumulated. Consequently, PCR bias can result in high error rates and missing population in NGS. For error-free data recovery, Erlich and Zielinski increased the NGS coverage from 10.4 (before multiple PCR reactions) to 69.4. At an increased NGS coverage, they were able to obtain the population with low abundance. In summary, the PCR method requires increased NGS coverage to recover the data, and this will increase the cost and duration of the data access process of DNA-based data storage. If the user uses the same NGS coverage to generate the DNA library using multiple PCR reaction, the user will fail to recover the data (Figure 3 of the manuscript).

Hence, the universal condition for the data access, including NGS coverage, cannot be determined using PCR-based methods. The data storage system should inform the user about the required NGS metrics for data access, according to the number of PCR reactions applied on the DNA library, by performing the preliminary experiment. Conversely, the user should apply excessive NGS coverage for every data access trial. However, this approach will consume cost and time in terms of data access in DNA-based data storage.

Also, Erlich and Zielinski[4] amplified and accessed the whole files simultaneously in the DNA library. However, because accessing whole files in the library is both time-

and cost-intensive, more recent studies have adopted the random access method, which enriches DNA molecules of a specific file from the DNA library using a DNA barcode to selectively access the data[5]. Multiple data accessing using the random access approach can be performed by applying multiple reactions of PCR by designing a universal primer for all molecules. Nevertheless, error rate according to PCR bias in the whole DNA library population can be predicted based on preliminary experiments. However, the abundance of molecules according to each file in the DNA library is random and unpredictable because of PCR stochasticity[14] (or randomness of amplification yield of each molecule during the early cycles of PCR). As a result, random access systems that apply PCR cannot determine the extent of NGS required for selective file recovery. In summary, PCR-based multiple accesses require change of data access condition and the time and cost for data access must be increased as the data is read. The consistency in data access condition is important in conventional data storage systems, such as hard disks.

When it comes to the physical density, Erlich and Zielenski[4]'s method should store multiple copies of DNA libraries for reamplification after the data read are stored. Their reading process is as follows; aliquoting 30 or 25 copies, amplifying one copy, and restoring amplified DNA libraries that remain after the data reading. Therefore, after two reactions, hundreds of copies of DNA library (30x25) must be stored and this lowers the physical density by two orders. As a result, recursive PCR method will lose physical density, more than $10^{12}$-fold after 9 reactions of recursive PCR. Physical density is one of the huge merits of DNA-based data storage to inspire general readership to imagine the DNA as the next-generation storage method. In contrast to this, DNA micro-disk utilize the original copy without alteration for multiple data reading of data, preserving the physical density.

ii) Hybridisation capture method

Hybridisation capture could be utilized to selectively enrich the specific file from the file library of random access to enable multiple accesses. Usually, complementary probes of 60mers or more are generally used for the hybridisation capture[12]. However, since the general index length of random access of 40mer is shorter than the length of typical probes, the capture sensitivity will be lower. Increasing the length of the probe in current DNA synthesis technology will increase the cost since the cost for synthesis is the major cost source for DNA-based data storage[6]. Additionally, due to the DNA synthesis length limit (~200nt), longer probe length will reduce information capacity (amount of the data stored in a DNA molecule) by occupying around 30% of DNA length and lower the physical density also. If we assume that the length of the index is long when the current state-of-art technology in hybridisation is applied, the specificity of hybridisation can be a problem. The current exome hybridisation method capture rate has a 20% off-target rate while capturing 30 ~ 50 megabases in 3 gigabases of an exome. The off-target ratio is normally inversely proportional to the target size, and if the target is small at several tens of kilo bases, such as BRCA 1 and 2, about an 80% off-target rate occurs[12]. From this off-target rate, the selection process without 100% specificity will corrupt the library of multiple files by capturing un-selected files in an unpredicted manner.

**Supplementary Note 2: Physical density**

For the physical densities, the redundancy of the same DNA molecule and the actual information capacity reduction due to the data in the DNA encoding design should be reflected. As a result, it was calculated as:

$$Physical\ density\ = \frac{DNA\ concentration\ *\ nucleotide\ per\ molecule\ (nt/molecule) * net\ information\ capacity(bit/nt)}{DNA\ molecule\ redundancy}$$

In general, the physical density is known to fall by about a ten thousand-fold compared to the ideal limit[23]. Using the encoding method that we previously introduced[6], we could recover information (image file of Hunminjeongum-Hyerye, Supplementary Fig. 8) without error when the information capacity of 1.27 and redundancy of 500 were used. The physical density was reduced by approximately 760 times compared to the limit (Fig. 2d). This result is similar to the data previously introduced. We believe that this value can reach the limit through the future technological development of DNA-based data storage methods.

In our experiment, the maximum capacity of the 160-bp double stranded DNA on the disk was approximately $10^1$ pmol/mm$^3$. The approximate molecular weight (g) of double stranded DNA is: number of nucleotides × 607.4 + 157.9; the mass capacity of DNA on the disk is approximately 973 ng/mm$^3$. We believe that physical density can be increased by advances in the DNA-based data storage field, including encoding and decoding techniques. DNA micro-disk is a carrier system of data-encoding DNAs; the application of the advanced technique will not be an issue. For example, the physical density of the DNA micro-disk can be increased to $1.32 \times 10^{14}$ bit/mm$^3$ by applying the state-of-the-art technique of DNA-based data storage that can store 17 exabytes/g (or $2.31 \times 10^{17}$ bit/mm$^3$) [22] (Fig. 2d).

**Supplementary Note 3: Acceleration test of DNA micro-disk**

We designed an experiment to examine the DNA integrity by checking the stability of the dried DNA pellet, DNA micro-disk in 50% relative humidity at various temperatures of 60, 65, and 70°C for two weeks. The 50% humidity level was controlled by placing a saturated salt solution (sodium bromide, Sigma Aldrich) in a gas-tight container. Then, the micro-disk containing ~$5x10^8$ DNA molecules or DNA of $6x10^{10}$ molecules was freeze-dried in a laboratory tube and placed in the container. After storage, the DNA micro-disk was washed three times with pure ethanol and used for DNA production. The quantity of DNA was validated via qPCR (Supplementary Fig. 3). By fitting the results with a first-order decay rate expression, we acquired the decay rate constants (*ln*(C) vs. time, C is the ratio of DNA when compared to the initial value. Supplementary Table 1). Additionally, we were able to acquire the Arrhenius-type activation energies of the Arrhenius equation following linearly fitting of *ln*(k) vs. 1/T. The value was within a range from 145 to 148 kJ/mol, which was aligned with the experimental values from previous reports of 120-155 kJ/mol (Supplementary Table 2). Additionally, after 2 weeks of storage, the code and shape of the disk was also maintained after several washes, enabling data management (Supplementary Fig. 4). Finally, we extrapolated the half-life of storage of the DNA micro-disk in 50% relative humidity as a function of temperature, which has been added in the manuscript in Fig. 2e. At temperatures lower than 10°C, the half-life of storage will be higher than 100 years (>700 years at 0°C).

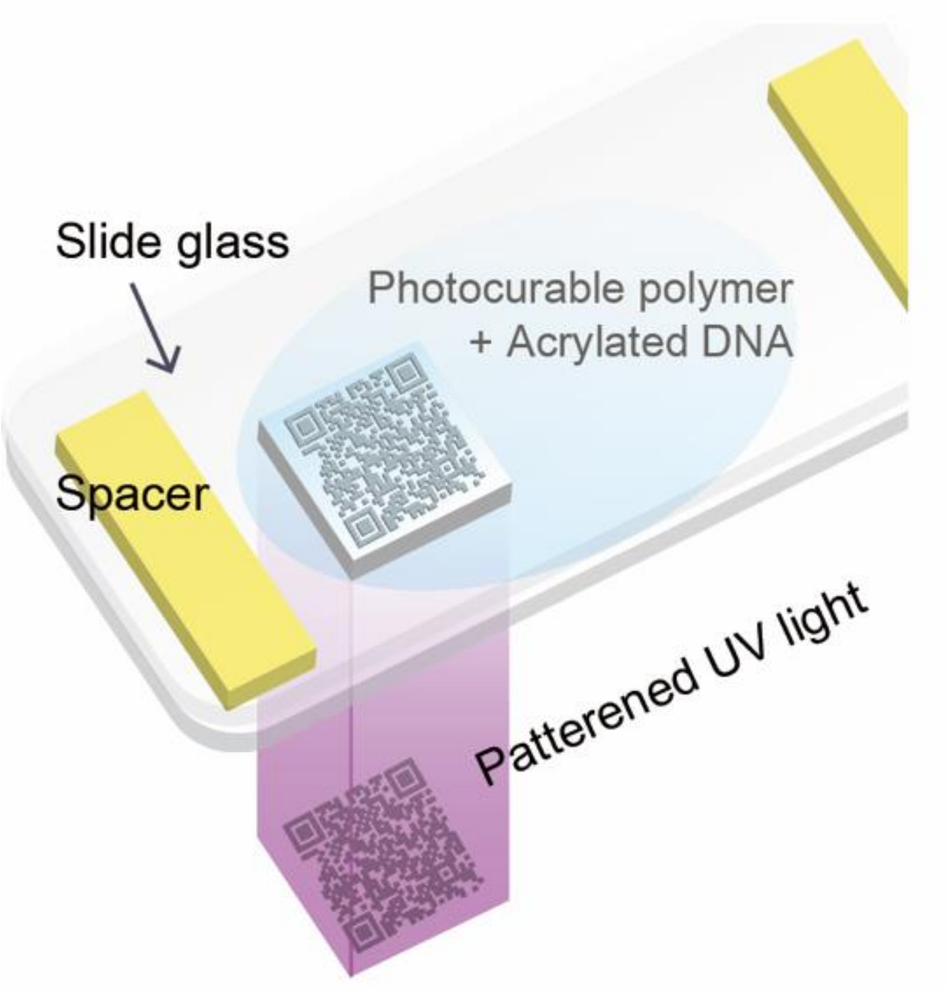


**Supplementary Figure 1**

DNA micro-disks are fabricated by OFML (optofluidic maskless lithography)[16].

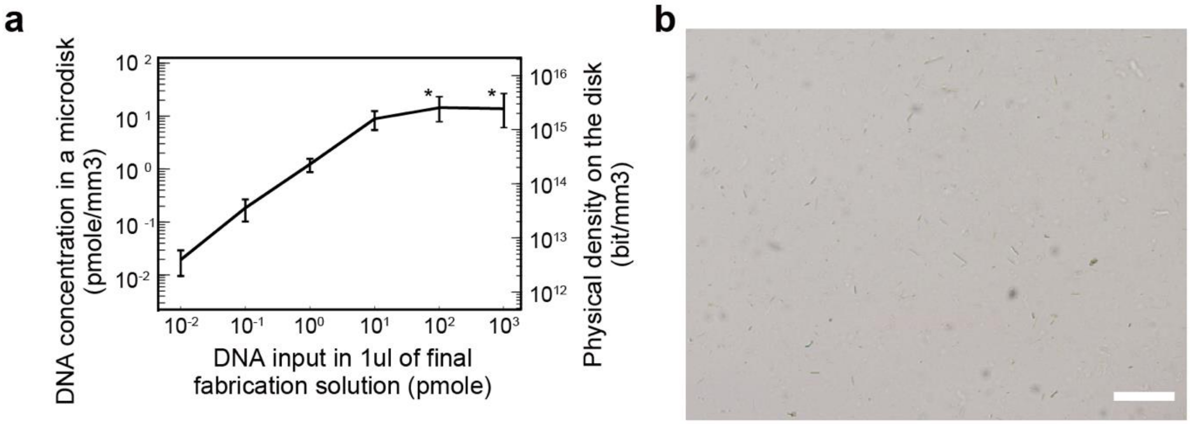


**Supplementary Figure 2**

**a**, Depending on the DNA concentration in the pre-polymer solution during fabrication, a micro-disk can be fabricated with various data capacities (n= 10). When the input DNA is more than 100 pico moles in 1 µl of solution, the DNA is not fully dissociated in the fabrication solution and is precipitated (indicated by asterisks, Supplementary Fig. 2b). The information capacity limit describes the maximum data amount that could be stored in ideal DNA-based data storage. Details about the calculation method are described in the Supplementary Note 2.

**b**, Precipitation of DNA in the polymeric resin is observed. Scale bar: 500 µm.

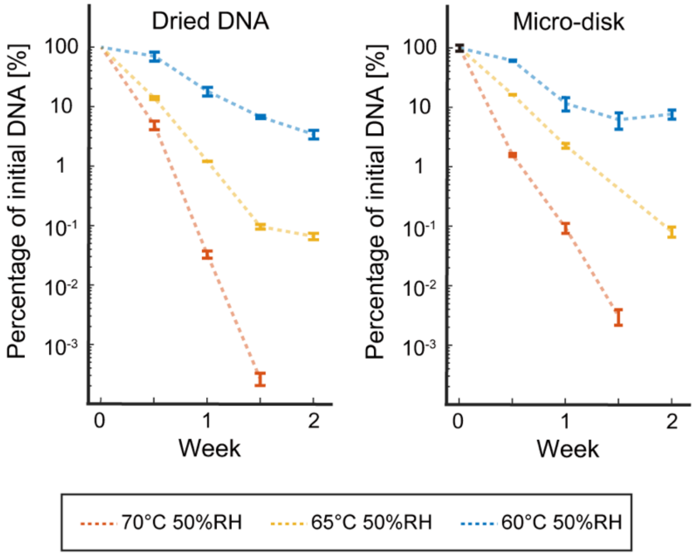


**Supplementary Figure 3**

Effect of the storage conditions on the DNA integrity. DNA integrity in a micro-disk was measured by qPCR, at various temperatures of 60, 65, and 70°C with 50% relative humidity for two weeks. The error bars represent the standard deviations (n=10).

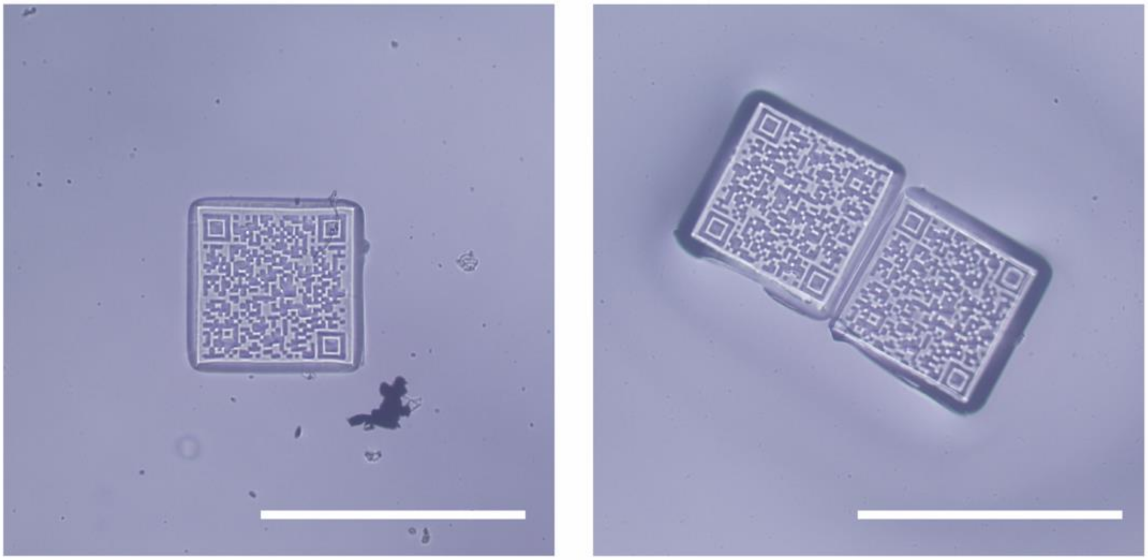

**Supplementary Figure 4**

DNA micro-disks after 2 weeks stored at 70°C with 50% relative humidity. Scale bar: 500 µm.

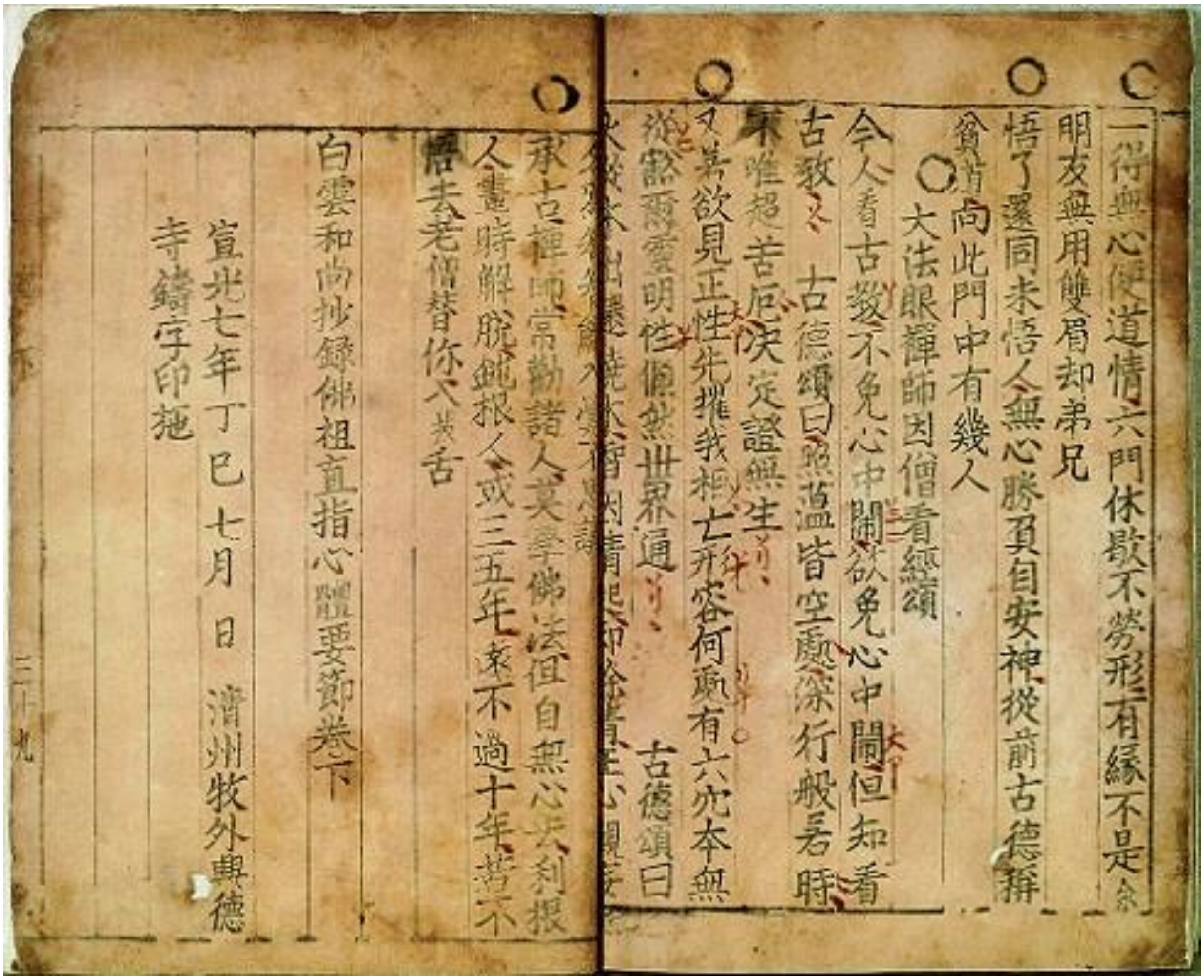

白雲和尚抄錄佛祖直指心體要節卷下

宣光七年丁巳七月 日 淸州牧外興德寺鑄字印施

**Supplementary Figure 5**

The thumbnail image of Jikji. Korean Buddhist document. Jikji is the world's oldest metalloid type. (https://en.wikipedia.org/wiki/Jikji)

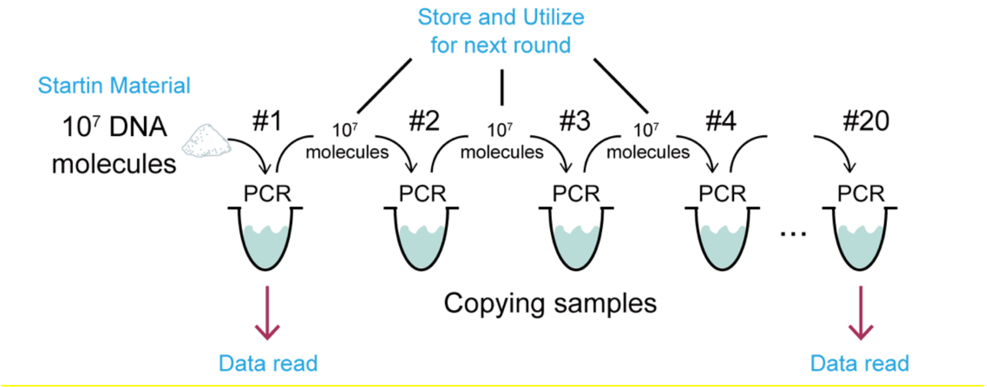


**Supplementary Figure 6**

In conventional DNA-based data storage, the data can be re-read by amplifying the original DNA molecules and re-amplifying the amplified molecules.

For the comparison of data loss, we set physical density as a fixed parameter for both methods as mentioned above. During the recursive PCR reactions, we amplified DNA molecules, read the data, and stored same amount of DNA molecules from PCR amplicon for next reaction of PCR, to preserve the physical density.

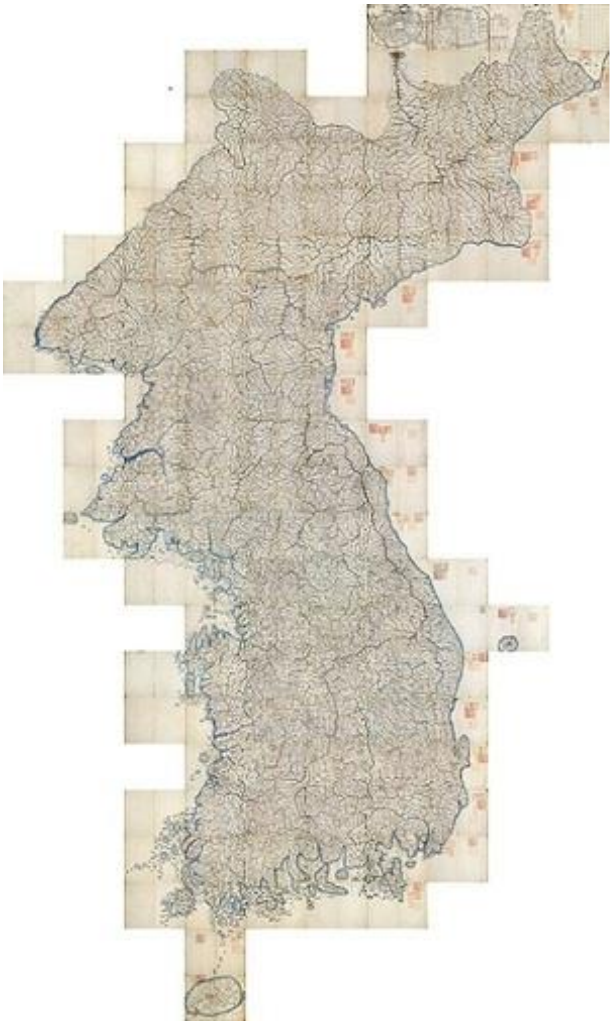

**Supplementary Figure 7**

The thumbnail image of Daedonyeojido, a map of Korea produced by Kim Jeong-ho in 1861, Joseon dynasty. (https://en.wikipedia.org/wiki/Daedongyeojido)

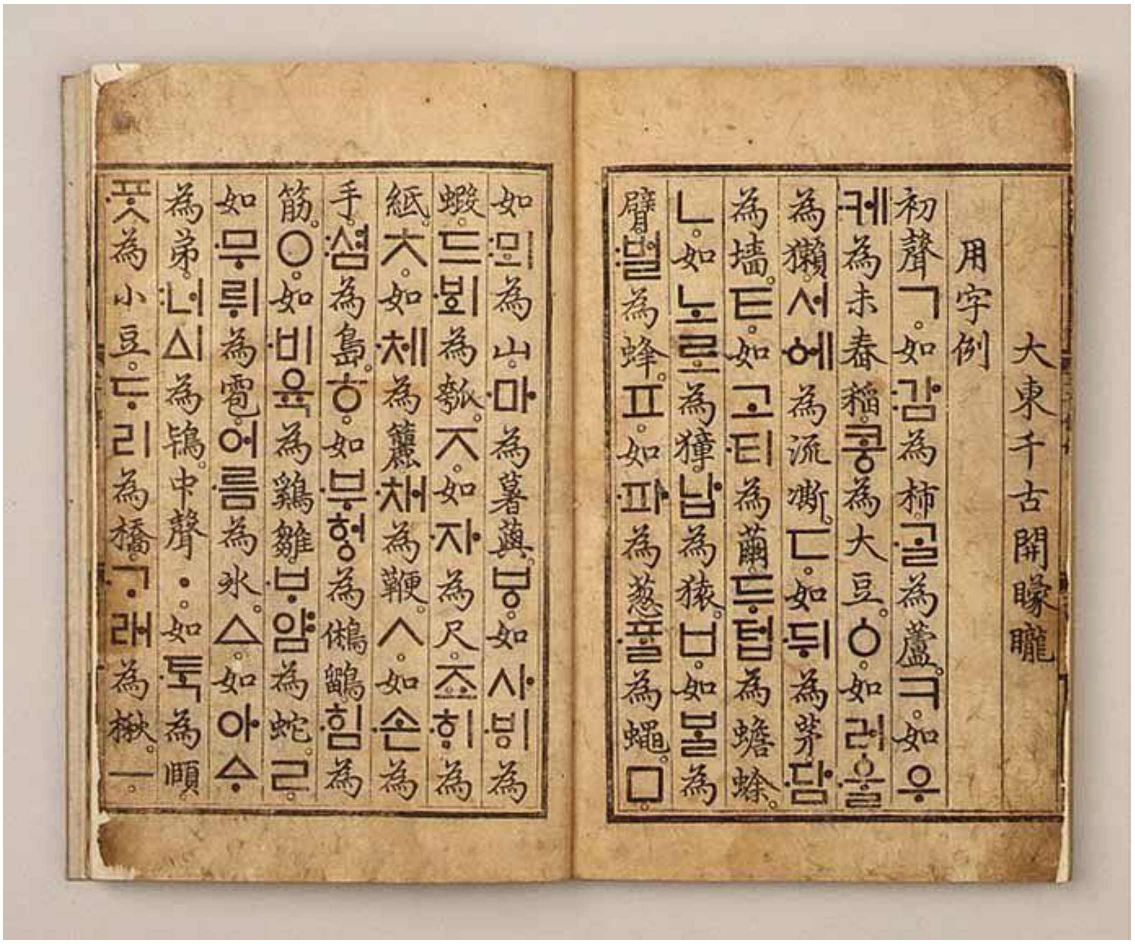


**Supplementary Figure 8**

The thumbnail image of Hunminjeongum Manuscript (or Hunminjeongum Haerye), which is the UNESCO memory of the world registered documented heritage submitted by Republic of Korea in 1997. This image was originally posted by the Cultural Heritage Administration of the Republic of Korea under the Korea Open Government License (https://www.mcst.go.kr/kor/s_open/kogl/koglType.jsp?pTab=05, The user can freely use the public work regardless of its commercial use without fee, and can change it to create secondary work.) type 1.

**Source:**
http://heritage.go.kr/heri/cul/culSelectDetail.do?region=1&searchCondition=&searchCondition2=&s_kdcd=11&s_ctcd=00&ccbaKdcd=11&ccbaAsno=00700000&ccbaCtcd=11&ccbaCpno=1111100700000&ccbaCndt=&stCcbaAsno=70&endCcbaAsno=70&stCcbaAs

dt=&endCcbaAsdt=&ccbaPcd1=99&culPageNo=1&chGubun=&header=view&returnUrl=%2Fheri%2Fcul%2FculSelectViewList.do&sCond=
