## Supplementary material for "DNA micro-disk for the management of DNA-based data storage with index and write-once-read-many (WORM) memory features": Materials and Method

### Materials and Methods

#### Encoding and decoding of DNA-based data storage

The DNA-based data storage encoding was performed as described in previous studies[6,11]. Briefly, binary data were extracted from the file and transformed by DNA codons. Then, the DNA sequence was fragmentised, and inner and outer Reed-Solomon error correction blocks were attached to the fragments. Finally, the address in the DNA codon form, forward primer, and reverse primer were attached. For the physical density experiments in Fig. 2d, 135 KB of Hunminjeongum was encoded into 160nt 5299 DNA fragments. For other experiments, 64 KB of the Jikji file was encoded into 143nt 5173 DNA fragments, 81 KB of Hunminjeongum was encoded into 143nt 6532 DNA fragments, and 55KB of Daedonyeojido was encoded into 143nt 4355 DNA fragments. The designed sequences are described in the Supplementary Files 1-4. For decoding, pair-end reads of the raw Fastq file were stitched using PEAR. Subsequently, sequencing read with encoded DNA length were filtered. Filtered reads were categorised by address and transformed to bits. Finally, error correction using Reed-Solomon was performed. For quality analysis (results shown in Fig. 3b), the reads were aligned to the design using the mem aligner of BWA (http://sourceforge.net/projects/bio-bwa/files/), followed by processing with SAMtools: view, sort, and mpileup.

#### DNA synthesis and sequencing

DNA for the data storage was synthesised via microarray-derived DNA oligopool synthesis. The B3 Synthesizer DNA microarray synthesizer (Customarray Inc.) was used. A 12k microarray was synthesised following a standard protocol. For sequencing, the DNA product, concentrated in 1 μL, was first amplified using the designed primer by qPCR (FAST 7500, Applied Biosystems) and the KAPA HiFi Library Amplification Kit. A sample mix of 10 μL master mix, 6 μL of PCR grade water, 1 μL each of a 10 μM stock of Forward and Reverse primers, and 1 μL of 20X SYBR Green was used. A standard thermal protocol was performed according to the manufacturer's instructions. Further, the amplification plot was assessed using qPCR. Once the plot reached saturation, the reaction was stopped, and the sample was purified using the PCR purification kit (Qiagen). Finally, the purified product was sequenced on a MiniSeq or HiSeq using a 300-cycle pair-end read protocol.

**DNA quantification**

Samples were analysed by qPCR (FAST 7500, Applied Biosystems) using a KAPA SYBR® FAST qPCR Master Mix (2X) Kit. Briefly, a sample mix of 10 μL master mix, 7 μL of PCR grade water, 1 μL each of a 10 μM stock of forward and reverse primers, and 1 μL sample was used. A standard thermal protocol was performed according to the manufacturer's instructions. Relative sample quantification was conducted by interpolation from a standard curve, generated from DNA samples of known concentration.

**Primer design**

For the physical density experiment in Fig. 2, the following primer sequence for the data-encoded DNA was used: Forward: (Acrydite) ACACGACGCTCTTCCGATCT and Reverse: GACGTGTGCTCTTCCGATCT. For experiments in Figs. 2 and 3, the following primer sequence was used: Forward: (Acrydite) ACACTCTTTCCCTACACGACGCTCTTCCGATCT and Reverse: GTGACTGGAGTTCAGACGTGTGCTCTTCCGATCT. For both designs, the final length of the amplicon was 160 bp after PCR processing. Additionally, the reverse primer was used for *in situ* DNA production. For the experiment shown in Fig. 4, the sequence of the forward primer was the same, and the TruSeq adaptor with i7 index of 707, 708, and 709 was attached to the reverse primer of the second experiment for each data-encoded DNA set. Primer CAAGCAGAAGACGGCATACGAGAT was used for *in situ* DNA production. For random access, Forward: ACACTCTTTCCCTACACGACGCTCTTCCGATCT and Reverse: CAAGCAGAAGACGGCATACGAGAT[i7] were used for amplification. All primers were ordered from the Macrogen.

**Maskless lithography of the micro-disk**

The prepolymer resin was prepared as a mixture of 41 vol% of poly(ethylene glycol) diacrylate (PEGDA, Mn~700, Sigma Aldrich), 41 vol% of poly(ethylene glycol) (PEG, Mn~600, Sigma Aldrich), and 9 vol% of 3-(trimethoxysilyl)propyl acrylate (TMASPA, Sigma Aldrich) as a alkoxysilane-grafted photocurable resin with 9 vol% of Irgacure® 1173 (BASF) as a photoinitiator. Various concentrations of data-encoded DNA library were prepared through amplification with a primer set, one of which was acrydite-modified. The purified DNA solution in water, after the amplification, was then mixed with the prepolymer resin at a 1:9 ratio by vigorously vortexing for 5 min at 2500 rpm. An optical microscope (IX71, Olympus), ultraviolet (UV) light source (365 nm, Lightningcure LC8, Hg-Xe lamp, Hamamatsu), and digital micromirror device (DMD, Texas Instruments) were aligned to modulate the UV light to a designed QR code pattern using the DMD pattern. Finally, the DNA micro-disk was fabricated by projecting the QR-patterned UV light to the dispensed resin between the polydimethylsiloxane (PDMS)-coated glass slides with proper spacer materials through a 10x (NA 0.3) objective lens. The micro-disk was then stored in freeze-dried conditions, for further use or storage.

**Assembly of the micro-disk**

Micro-disks were assembled on the PDMS well by self-assembly following previous studies[20] or directed assembly using tweezer on the PDMS well. For the PDMS well plate, the conventional soft lithography process was used. The master mould for the PDMS well was fabricated using a negative photoresist SU-8 (Microchem Corp.) with the lithography process. The polydimethylsiloxane (PDMS, Sylgard 184, Dow Corning) copolymer was mixed with a curing agent (10:1 w/w) and poured on the master mould. After removing micro bubbles, the PDMS well plate was baked at 150°C for 10 min and removed from the mould.

***In situ* DNA production cycle and multiple reading experiment**

DNA micro-disk was selected, transferred to 20 μL of 0.15 N Sodium hydroxide (NaOH, Sigma Aldrich) in a laboratory tube, and incubated for 5 min for denaturation. Following, the supernatant was collected, and the pH was adjusted to neutral by adding 10x Tris-EDTA (TE, Sigma Aldrich) and Acetic Acid (Sigma Aldrich), for used as a DNA copy from the disk. The micro-disk was washed in 1x Tris-EDTA five times, 70% ethanol once, and dried. Subsequently, the primer was hybridised, and elongation proceeded by adding 25 µL solution containing 0.5 µL of Bst 2.0 WarmStart® DNA Polymerase (New England Biolabs), 0.5 µL of 10 mM dNTPs (New England Biolabs), 0.5 µL of 10 µM primer, and 2.5 µL of 10x standard reaction buffer. The reaction solution was incubated for 10 min at 55°C. The micro-disk, which contained double-stranded DNA on its polymeric mesh, was washed in 1x Tris-EDTA five times, 70% ethanol once and freeze-dried for further use. For the experiments in conventional multiple readings of DNA-based data storage (Fig. 3b, Supplementary Fig.6), $10^7$ molecules of DNA, concentrated in 1 µL, was amplified using the designed primer by PCR. A sample mix of 10 µL master mix, 6 µL of PCR grade water, and 1 µL each of a 10 µM stock of Forward and Reverse primers, was used. A standard thermal protocol of 17 cycles of PCR was performed according to the manufacturer's instructions. After the PCR, the sample was purified using the PCR purification kit (Qiagen), quantified, and used for the reamplification.
